# DYNATE: Localizing Rare-Variant Association Regions via Multiple Testing Embedded in an Aggregation Tree

**DOI:** 10.1101/2022.12.01.518706

**Authors:** Xuechan Li, John Pura, Andrew Allen, Kouros Owzar, Jianfeng Lu, Matthew Harms, Jichun Xie

## Abstract

Rare variants (RVs) genetic association studies enable researchers to uncover the variation in phenotypic traits left unexplained by common variation. Traditional single-variant analysis lacks power; thus, researchers have developed various methods to aggregate the effects of RVs across genomic regions to study their collective impact. Some existing methods utilize a static delineation of genomic regions, often resulting in suboptimal effect aggregation, as neutral subregions within the test region will result in an attenuation of signal. Other methods use varying windows to search for signals but often result in long regions containing many neutral RVs. To pinpoint short genomic regions enriched for disease-associated RVs, we developed a novel method, DYNATE (DYNamic Aggregation TEsting). DYNATE dynamically and hierarchically aggregates smaller genomic regions into larger ones and performs multiple testing for disease associations with controlled weighted false discovery rate. Extensive numerical simulations demonstrate the superior performance of DYNATE under various scenarios compared to existing methods. Importantly, DYNATE-identified regions have higher enrichment levels of disease-associated RVs. We applied DYNATE to an amyotrophic lateral sclerosis (ALS) study and identified a new gene, *EPG5*, harboringpossibly pathogenic mutations.

## 1 Introduction

Rare variant (RV) disease association analysis has been an important topic in genetics and genomics. Because single-variant analysis suffers low statistical power (Bansal et al., 2010; Lee et al., 2014), a common approach to increase statistical power is to test the collective effect of all RVs within genomic prefixed regions. The testing methods include burden tests (Li and Leal, 2008; Asimit et al., 2012; Morris and Zeggini, 2010), variant component tests (Neale et al., 2011; Wu et al., 2011), and their combinations (Lee et al., 2012). The prespecified regions could be genes (Petrovski et al., 2017; Povysil et al., 2019), functional domains (Gelfman et al., 2019), or sliding fixed-length windows (Bhatia et al., 2010; Katsumata and Fardo, 2020). However, those prefixed-region-based methods are suboptimal when the pre-fixed regions are too long or short. For example, when the pre-fixed region is too long, it could harbor too many neutral variants. These neutral variants will attenuate the effect of any causal variants in the region. On the other hand, when the pre-fixed region is too short, even if the region contains some causal variants, the analysis may exclude adjacent regions containing additional causal RVs, leading to a loss of power. An extreme example of the latter is the single-variant association analysis.

Inspired by these considerations, varying-window methods (Li et al., 2019; Kanoungi et al., 2020; Li et al., 2022b,a; He et al., 2019) scan subregions with different lengths to optimize the power of detecting disease-associated RVs. Compared with fixed-region methods, varying-window methods are generally more powerful. However, in practice, the regions identified by the varying-window methods often include long stretches of sequence containing many neutral RVs; thus, they are less helpful in pinpointing the functionally important RVs in the downstream analysis. Moreover, searching regions with different lengths is computationally intensive, leading to long computing times.

This paper proposes a new method that pinpoints short regions containing enriched disease-associated RVs. The method, called DYNATE (DYNamic Aggregation TEsting), dynamically aggregates and tests genomic regions with varying lengths for disease association. DYNATE adopts a fine-to-coarse hierarchical testing strategy, so that, given the previous layer’s testing results, DYNATE dynamically defines the testing regions, performs multiple testing on the current layer, and passes the results to the next layer. The hierarchical testing structure allows DYNATE to explore both short and long regions efficiently, resulting in a fast computational algorithm.

We demonstrated that DYNATE outperforms other competing methods under various numerical settings: DYNATE has similar power as the state-of-the art fixed-region (Gelfman et al., 2019) and varying-window (Li et al., 2019) methods in identifying disease-associated domains. However, its RV-level precision is much higher: the DYNATE-identified regions are shorter and highly enriched with pathogenic variants, while the competing methods often output extensive regions that cover too many neutral variants. We applied DYNATE to an amyotrophic lateral sclerosis (ALS) study to pinpoint the regions containing ALS-associated RVs. DYNATE successfully identified regions in well-known ALS genes *SOD1* (MIM: 147450) and *TARDBP* (MIM: 605078) while also identifying a new region in *EPG5* (MIM: 615068), which could not be identified by competing methods. Further, we compared the identified region lengths. The DYNATE-identified regions were much shorter, suggesting a higher disease-associated RV specificity. Thus, DYNATE provides more specific targets for the downstream functional pathway analysis.

## 2 Materials and Methods

### 2.1 Notations

DYNATE adopts a dynamic, hierarchical testing framework that decomposes the whole-exome or whole-genome into smaller segments, referred to as leaves. These leaves are subsequently consolidated into larger segments for testing purposes. The notations utilized in this paper are outlined below.

#### 2.1.1 Leaves and nodes

Assume a case-control study with *N*_0_ controls, *N*_1_ cases, and *J* observed qualifying RVs (Rare Variants) in *D* homology-based protein domains (Lawrence and Goldman, 1988). DYNATE operates on various layers represented by superscript ℓ.

In the first layer (layer 1), these *D* homology-based protein domains are fragmented into *m*^(1)^ leaves to amplify resolution and identify disease-associated RVs. A *leaf* is described as a genic region where *M* subjects present with continuous qualifying rare variants. Here, *M* denotes leaf size, which is significantly smaller compared to *N*_0_ and *N*_1_. When *M* = 1, each leaf possesses qualifying RVs from a single subject. As a general rule, we suggest *M ≥* 5.

Applying protein domain annotations, we further refine leaf definition by enforcing the disintegration of leaves at domain extremities. Consequently, a leaf does not extend across two domains. If fewer than *M* subjects present with qualifying variants in the final leaf of a domain, this leaf is merged with the preceding leaf in the same domain, if present. If a domain contains less than *M* qualifying variants, the entire domain region forms a leaf.

Assume the partition culminates in *m*^(1)^ leaves, represented by 1,…, *m*^(1)^. On superior layers (layer *ℓ* ≥ 2), it aggregates the accepting nodes from the preceding layer to formulate new nodes. A node *S* represents the union of one or multiple leaves, characterizing a genome region with qualifying RVs. Therefore, leaves are nodes on the first layer. For any node *S*, the set of its qualifying RVs is represented by 𝒱 (*S*), with 𝒱 (*S*) ⊂ [*J*].

On all layers, we categorize qualified nodes as those newly formulated on the current layer that have not been previously tested. On the first layer, all leaves qualify as nodes. On layer *ℓ* (*ℓ* ≥ 2), a qualified node represents the union of two or more contiguous nodes on layer *ℓ -* 1. The set of all qualified nodes on layer *ℓ* is represented by *B*(^*ℓ*)^. Specifically, *B*^(1)^ = *i* : *i ∈* [*m*^(1)^], where [*m*^(1)^] represents the set 1,…, *m*^(1)^.

#### 2.1.2 Node hypotheses and P-values

We call leaf *{i}* a null leaf if for all *j 2 ∈ 𝒱* (*{i}*), the qualified RV *j* is not associated with the disease; otherwise, we call it alternative. Denote the set of null leaves by 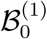 and the set of alternative leaves by 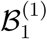.

Similarly, for any node *S*, we denote the node hypothesis by *H*_*S*_. The null is

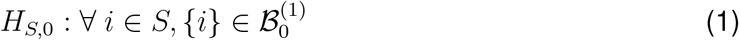

For simplicity, for leaf hypothesis *H*_*{i}*_, we denote them as *H*_*i*_. *H*_*i*_ is null if *{i} ∈* 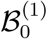 and alternative if *{i} ∈* 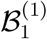.

To test these hypotheses, we need to summarize their P-values. DYNATE is flexible to accommodate existing RV test statistics including burden tests(Asimit et al., 2012; Morgenthaler and Thilly, 2007; Li and Leal, 2008), variance-component tests(Wu et al., 2011; Pan, 2009), and omnibus tests(Lee et al., 2012). For fast computation, we mainly used the following two statistics: 1) Lancaster’s mid-P correction for the Fisher’s exact test (FL), and 2) efficient score statistics with saddle point approximation (SS). More details are provided in Supplementary Note 1.

After leaf P-values are calculated, we use Stouffer’s method to calculate node P-values. For node *S*, the P-value is

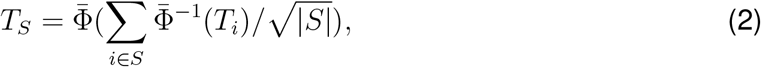

where 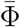(·) is the complementary cumulative density function of the standard Gaussian distribution. Alternatively, we may consider using other aggregation approaches, such as Fisher’s combination(Fisher, 1925), chi-square aggregation, or Cauchy aggregation(Liu and Xie, 2020).

#### 2.1.3 Type I Error criterion: node-FDR

When a node is the union of multiple leaves, we might not be able to dichotomize its null and alternative status. For example, suppose node *S* = *{*1, 2*}* with leaf *{*1*}* is null and leaf *{*2*}* is alternative. Then *S* is ½ alternative. More generally, for any node *S*, let *S*_1_ = *S ∩ {i* : *H*_*i*_ is alternative*}*. Then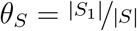. We refer to *S* as θ_*S*_-alternative or (1 *θ*_*S*_)-null.

To address this fractional null/alternative status of nodes, we introduce a new type I error measure called node-false-discovery-rate (node-FDR), which is the expectation of node-falsediscovery-proportion (node-FDP)

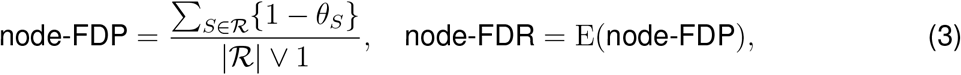

where *R* is the set of the rejected nodes. Rejecting node *S* will contribute 1 *θ*_*S*_ to the numerator and 1 to the denominator.

When identifying disease-associated regions, DYNATE aims to asymptotically control node-FDR. Thus, it prioritizes identifying the nodes containing higher proportions of alternative leaves. For example, when the alternative leaves are concentrated in a shorter region, DYNATE will prioritize identifying this short region rather than a more extended region containing too many null leaves. Because these shorter regions usually contain a higher proportion of disease associated RVs, they are more likely to be pathogenic or protective.

### 2.2 DYNATE: Dynamic and Hierarchical Testing

To identify disease-susceptible regions, DYNATE adopts a fine-to-coarse strategy. First, it tests the leaf-level disease associations, where leaves are the smallest testing units. Then, it hierarchically aggregates leaves into nodes (corresponding to longer regions) and tests the node-level disease associations. The general algorithm is listed in Algorithm 1. To illustrate the implementation of Algorithm 1, we provided a toy example in Supplementary Figure S1.

#### Algorithm 1

Dynamic hierarchical testing

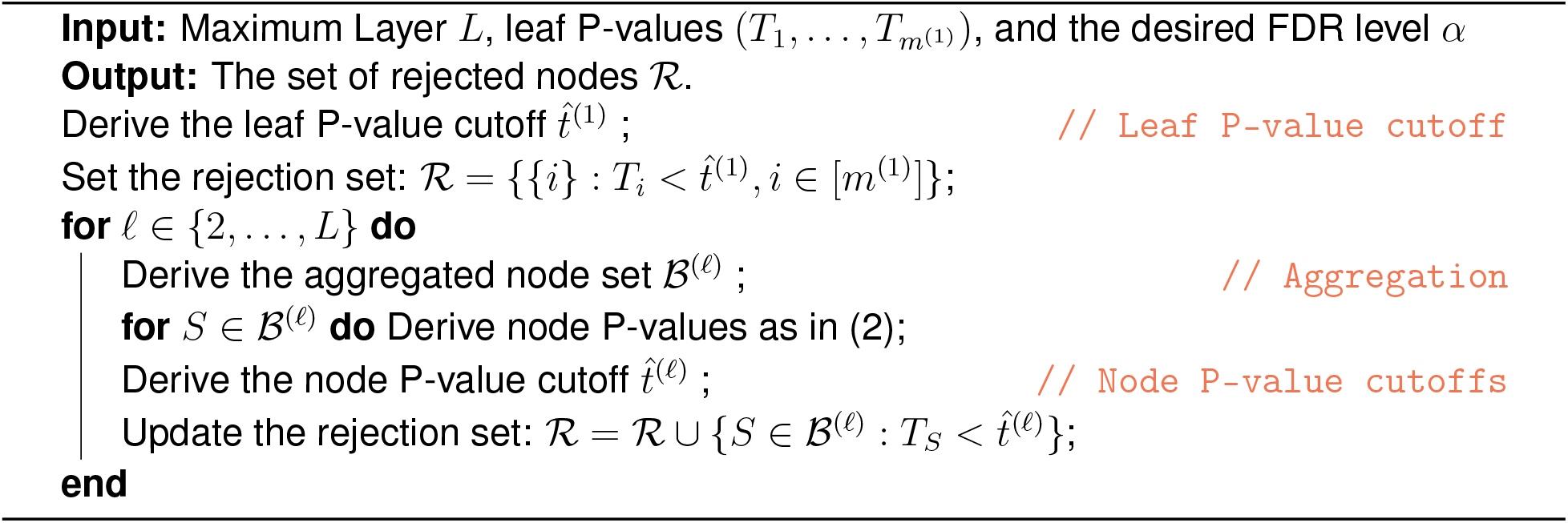

Algorithm 1 includes multiple steps: leaf P-value cutoff, aggregation, and node P-value cutoffs. Another key step is to choose tuning parameters. Here, we provide a summary of each step; the details were provided in Supplementary Note 2.

- *Leaf P-value cutoff*. This step is to choose a proper cutoff 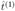 to control the node-FDR on layer 1. We noticed that on layer 1, the node-FDR is the same as the FDR for all leaf hypotheses; thus, using traditional FDR cutoff (such as the cutoff for the classic Benjamini and Hochberg procedure (Benjamini and Hochberg, 1995)) works well on layer 1.
- *Aggregation*. This step is to aggregate the accepted nodes on previous layers into larger nodes on the current layer. Here, “accepted” nodes refer to those not rejected on the previous layer. The purpose is to aggregate neighboring accepted nodes that are likely to be co-null or co-alternative; if they are co-alternative, aggregation will increase their chance of being rejected.
- *Node P-value cutoffs*. This step is to choose a proper cutoff 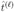 to control the node-FDR on layer *ℓ* with *ℓ* 2. Because a node can be partially null, we hope that we could roughly estimate nodes’ overall null level to provide a control of node-FDR. We use the sampling technique to get this overall estimate.
- *Tuning parameter selection*. The layer number *L* and leaf size *M* are tuning parameters. The default tuning parameters were set to optimize the RV-level specificity in the findings and the power. Our default setting was validated by the numerical studies shown in **Simulation Studies**.

### 2.3 Simulation Studies

We evaluated the performance of DYNATE under a wide range of simulation settings. We mimicked the structure of RVs and domains in the ALS dataset, which contains 263, 789 qualified variants located in 20, 666 qualified functional domains. Please see **Application to an Amyotrophic Lateral Sclerosis (ALS) Study** for how we pre-processed the data to obtain the qualified variants and domains. We randomly selected 2, 000 domains with 25, 622 RVs.

Out of the 2000 domains, we randomly selected 20 domains with more than 30 RVs to be dense pathogenic domains (DPDs) and 20 other domains to be sparse pathogenic domains (SPDs). A DPD is a domain harboring a pathogenetic region with min(20, *L*_*d*_*/*2) observed RVs, where *L*_*d*_ is the number of observed RVs in the domain. Within this pathogenic region, π of them are randomly selected as pathogenic RVs. In the simulation, we vary π *∈ {*20%, 40%, 60%, 80%*}* to evaluate the performance of DYNATE under different settings. An SPD is a domain harboring only one pathogenic RV; this RV location is randomly selected.

Let ***G***_*k*_ = (*G*_*k*,1_,…, *G*_*k,J*_)^*T*^ denote the observed RV vector for individual *k* where *G*_*k,j*_ is the indicator of whether individual *k* carries the mutation of RV *j*. We assumed that *G*_*k,j*_ *∼* Bern(ρ_*j*_) and simulated log_10_ *ρ*_*j*_ from Unif(*-*3, *-*1.7) so that *ρ*_*j*_ ranges from 0.001 to 0.02. Here, Bern(*ρ*) denote a Bernoulli distribution with parameter ρ, and Unif(*a, b*) denote a uniform distribution over the interval [*a, b*] respectively. When *ρ*_*j*_ is small, it is close to the minor allele frequency.

The binary phenotype of the individual *k* was independently generated based on the model

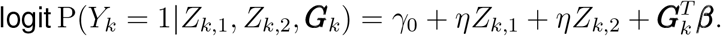

Here, *Z*_*k*,1_ and *Z*_*k*,2_ are two non-genetic covariates with *Z*_*k*,1_ ∼ *N* (0, 1) and *Z*_*k*,2_ *∼*Bern(0.5), respectively. Their coefficients are η, varying in different settings with *η ∈ {*0, 0.5*}*. The other set of coefficients β = (β_1_,…, β_*J*_)^*T*^ represents the RV effect size; we set β _*j*_ as a decreasing function of ρ_*j*_, β _*j*_ = *-c* log_10_ ρ_*j*_, where *c* = 2.5 for pathogenic RVs in the DPDs and *c* = 7.5 for those in the SPDs. We set the intercept, _0_ so that the disease prevalence is 1%.

Based on the 6 different combinations of π ∈ *{*20%, 40%, 60%*}* and η ∈ *{*0, 0.5*}* (covariate), we generated 6 large population, each with 2 million individuals. We conduct the numerical studies with 100 repetitions under each population. In each repetition, we randomly draw *N*_1_ cases and *N*_0_ controls. We let *N*_1_ = 1000 and vary *N*_0_ ∈ *{*1000, 2000, 3000*}* to check the performance of DYNATE when the number of controls increases.

When applying DYNATE to the simulated datasets, we used both the FL and SS statistics (See **Node hypotheses and P-values**). Under all scenarios, we set the layer number *L* = 3 and leaf size *M* = 11 according to the criteria described in **Tuning parameter selection**. In order to justify the default tuning parameters selection (Supplementary Note 2) and check the robustness of DYNATE, we also apply DYNATE with different *M* and *L* settings: *M ∈ {*7, 9, 11, 13*}*; for *M 2 {*7, 9, 11*}, L* = 3; for *M* = 13, *L* = 2.

We compared the performance of the DYNATE with competing methods, including the domain-based collapsing (DC) method(Gelfman et al., 2019) with the FL (DC-FL) and SS (DC-SS) statistics and the SGANG(Li et al., 2019) with their suggested burden (SCANG-B), SKAT (SCANG-S), and SKAT-O (SCANG-O) statistics and the default parameter setting. The desired domain-FDR for the DC method, familywise error rate (FWER) for SCANG, and node-FDR for the DYNATE are all set at 5%. The results yielded from these methods are evaluated using the average node-FDP, RV-FDP, RV-sensitivity, domain-FDP, and domain-sensitivity across repetitions. The node-FDP is defined in (3), and other measures are defined below:

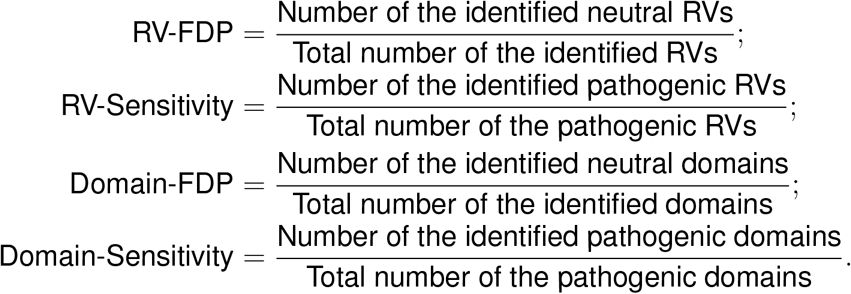

Here, an RV is called “identified” if it is located in one of the identified regions; a domain is called “identified” if any of its RVs, regions, or itself is called significant by the algorithms.

We also used RV-level and domain-level F1 scores as overall accuracy measurements: a high F1 score indicates high accuracy. The RV-level and domain-level F1 scores are defined below:

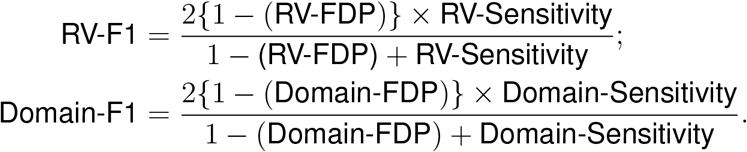

We used domain-level, node-level, and RV-level measurements to systematically evaluate the performance of DYNATE and the competing methods. Among them, DC(Gelfman et al., 2019) is a domain-collapsing method; it can only provide domain-level testing. Thus, the comparison between DYNATE and DC was conducted on the domain-level. SCANG(Li et al., 2019) is a varying-window scanning method; it can identify disease-associated regions with various lengths. Thus, the comparsion between DYNATE and SCANG was conducted on both region-level. Furthermore, we used the RV-level measurements to evaluate these methods’ abilities to pinpoint highly disease-susceptible genomic regions and narrow down the collection of possibly disease-associated RVs.

### 2.4 Application to an Amyotrophic Lateral Sclerosis (ALS) Study

We applied DYNATE to an ALS study with samples that have either whole-exome sequencing (WES) or whole-genome sequencing (WGS). The goal of this study is to pinpoint the pathogenic RV coding sequence regions in individuals with estimated European ancestry.

We started from the diversified population with 3239 cases and 11,808 controls. Qualifying variants were defined as nonsynonymous coding or canonical splice variants that have a minor allele frequency (MAF) ≤ 0.1% in cases and controls (internal MAF) and also a ≤ 0.1% MAF imposed for each population represented in the ExAC Browser(Lek et al., 2016). Gelfman et al. (Gelfman et al., 2019) filtered the variants based on their quality scores, quality by depth scores, genotype quality scores, read position rank sums, mapping quality scores, maximum Fisher’s strand bias, alternative allele ratios, et al. Variants were annotated to Ensembl 73 using SnpEff(Cingolani et al., 2012).

Based on the filtered variants, we defined a European sub-cohort from the diversified population. We required the individuals 1) to self-declare as “White”, 2) with an overall genotyping rate ≥ 0.87, and 3) with genotypes close to European, defined as the European ancestry probability ≥ 0.5. Here, the European ancestry probability was estimated through a multi-layer perceptron method used in Gelfman et al. (Gelfman et al., 2019).

The selected European sub-cohort contains 10, 138 samples: 2, 634 are cases (2, 590 samples from WES and 44 from WGS) and 7, 504 are controls (7, 239 samples from WES and 265 from WGS). There are 263, 789 qualified variants spanning 20, 666 qualified functional domains. The functional domains are defined as the genetic regions that aligned to the Conserved Domain Database (CDD)(Marchler-Bauer et al., 2012) and the unaligned genetic regions between CDD alignments(Gelfman et al., 2019). A functional domain is qualified if it contains more than 5 RVs.

We applied DYNATE with FL (DYNATE-FL) and SS (DYNATE-SS) statistics. The SS statistics were obtained from fitting model (S1) with no non-genetic covariates (η_*i*_ = **0**). The node-FDR level is 5%. We followed the tuning parameter selection criterion in Suplementary Note 2 to select the default leaf size *M* = 7 and layer number *L* = 5.

We also compared the results from DYNATE with those from SCANG and DC. For SCANG, we used their default tunning parameter settings and considered three types of statistics: burden (SCANG-B), SKAT (SCANG-S), and SKAT-O (SCANG-O). Following Li et al. (Li et al., 2019), the minimum and maximum numbers of variants in the sliding windows in SCANG were set as 1 and 103, respectively. For DC, we considered both FL (DC-FL) and SS (DC-SS) statistics. The desired FWER for SCANG and the domain-FDR for DC were set at 5%. A detected region was evaluated by its length and logOR. The region length is defined as the number of its containing RVs. The logOR is defined as

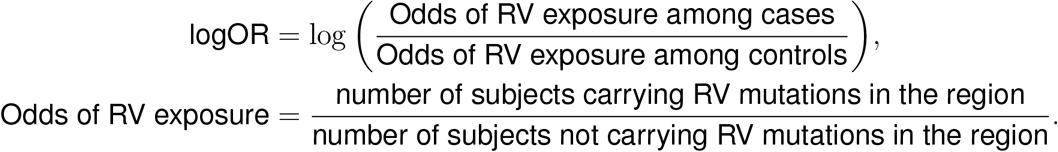

If logOR *>* 0, the region is likely to be pathogenic; if logOR *<* 0, the region is likely to be protective.

## 3. Results

### 3.1 Simulation Studies: Type I Error, Power, and Robustness of DYNATE

Figure 1 depicts the average node-FDP and RV-level sensitivity of DYNATE with the default tuning parameter setting. The node-FDP measures type I error and the RV-sensitivity measures power. We calculated the average node-FDR and RV-sensitivity at each layer up to the default layer number *L* = 3, with a 90% confidence interval to account for simulation variability. A slight inflation of the simulated node-FDR was observed, but it is important to highlight that the deviation is not significant. Therefore, it is still valid to claim that the simulated FDR remains closely aligned with the desired level.

**Figure 1:**
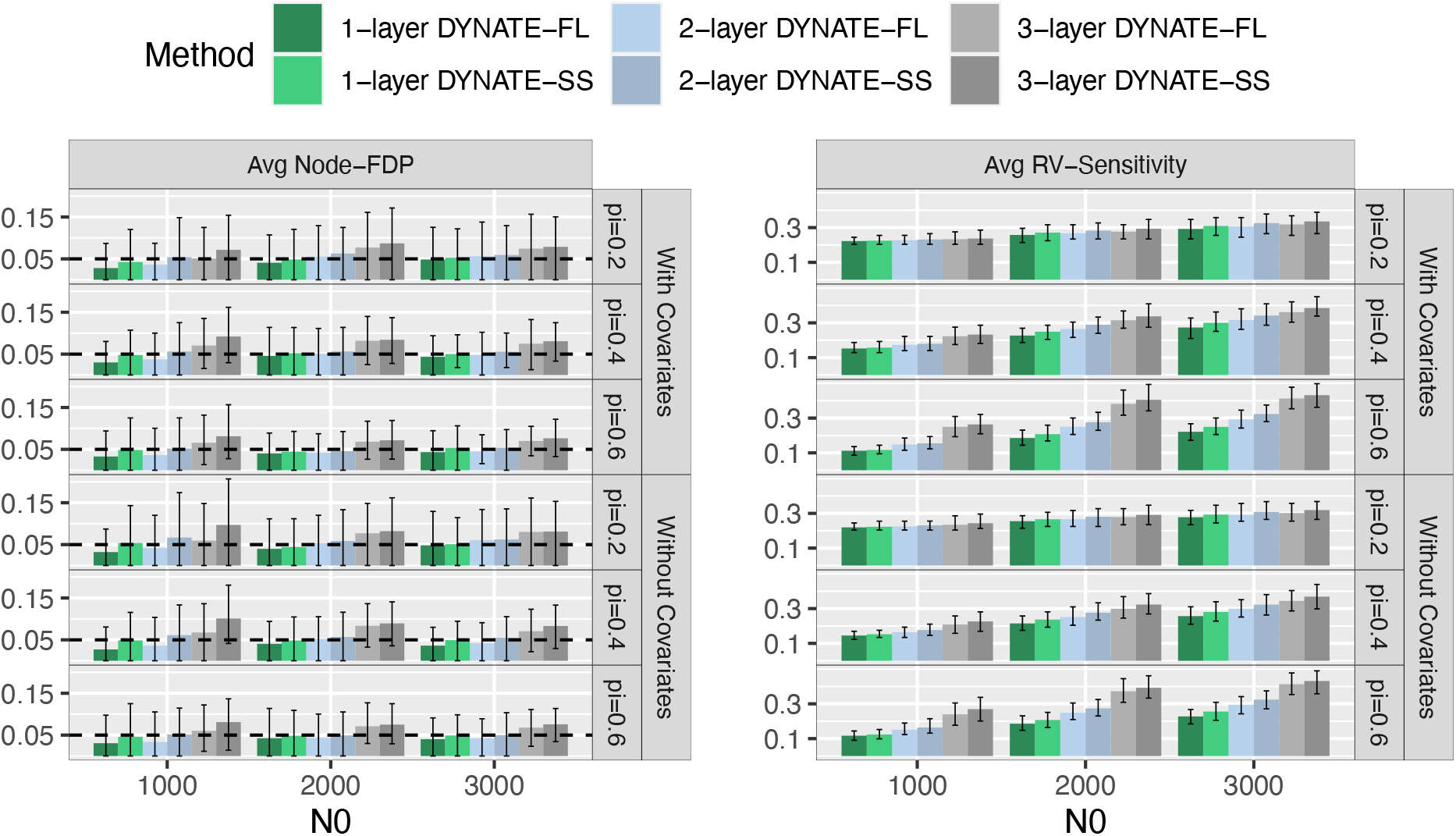
Performance of DYNATE after layer 1, layer 2, and layer 3. The number of cases *N*_1_ = 1000; the number of controls *N*_0_ ∈ {1000, 2000, 3000}. Different colors mark the DYNATE results with different statistics (FL and SS) and after different layers (1, 2, and 3). The main bars mark the average performance across 100 repetitions; the error bars mark the 90% confidence intervals – the 5% and 95% quantiles over the 100 simulations. The left panel shows the average node-FDP: the dashed horizontal lines mark the desired node-FDR level 5%. The right panel shows the average RV-sensitivity.

Meanwhile, the RV-sensitivity increases as the layer number goes up. The results indicate that the multiple-layer testing approach can significantly improve the analysis power. DYNATE with different choices of statistics (DYNATE-FL and DYNATE-SS) have similar performance under various settings. Sometimes, DYNATE-SS has a higher node-FDR, especially when the algorithm goes to layer 3, but DYNATE-SS also has higher RV-sensitivity than DYNATE-FL.

The robustness of DYNATE was further validated with varying leaf sizes *M ∈* 7, 9, 11, 13, as illustrated in Figure 2. Across all leaf sizes, the average node-FDPs of DYNATE ranged from 4% to 10%, with a target level of 5%. The default tuning parameter setting (*M* = 11 and *L* = 3, as established in **Tuning parameter selection**) provided a satisfactory average node-FDP (approximately 7%) and the highest RV-sensitivity. At *M* = 13, the algorithm halted at layer 2 due to a potential lack of remaining leaves to facilitate an additional layer of analysis. Although the absence of layer 3 enhanced average node-FDP control, it also impacted RV-sensitivity.

**Figure 2:**
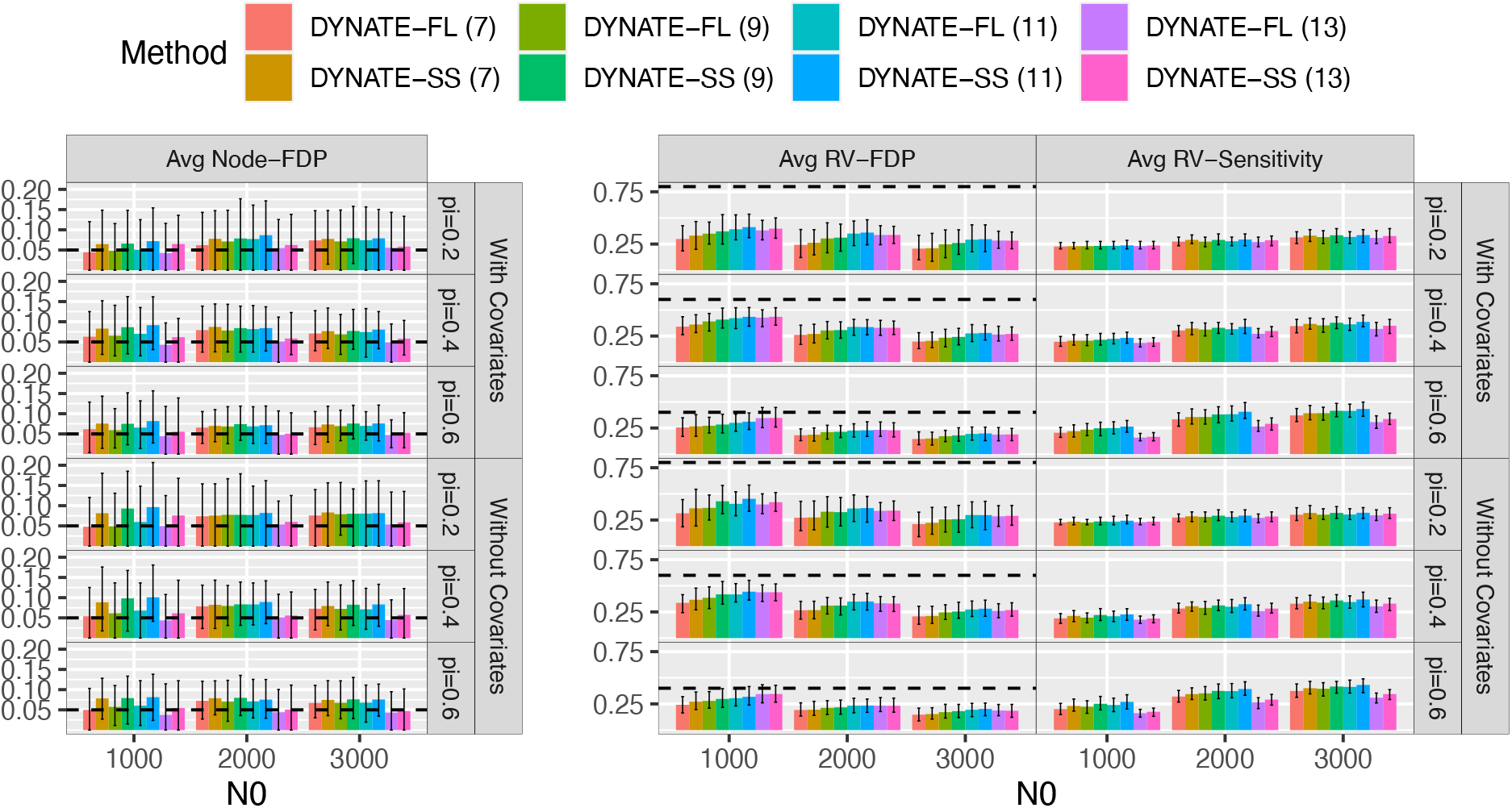
Performance of DYNATE across different leaf sizes. The number of cases *N*_1_ = 1000; the number of controls *N*_0_ ∈{1000, 2000, 3000}. Different colors mark the DYNATE results with different statistics (FL and SS) and different leave sizes (*M ∈*, 9, 11, 13 }). The main bars mark the average performance across 100 repetitions; the error bars mark the 90% confidence intervals – the 5% and 95% quantiles over the 100 simulations. The left panel shows the average node-FDP: the dashed horizontal lines mark the desired node-FDR level 5%. The middle panel shows the average RV-FDP: the dashed horizontal lines mark the neutral RV proportion in DPDs 1 *π*.. The right panel shows the average RV-sensitivity.

Figure 2 further revealed that DYNATE accurately identified numerous regions with enrichment of pathogenic RVs. Given our simulation setting, the proportions of neutral RV in the pathogenic region are 1 *π*. Therefore, the RV-FDP within the identified regions is anticipated to be less than 1 *π*.. Consistent results were observed with the RV-FDP in the DYNATE-identified regions. This evidence signifies the successful identification of subregions with significant pathogenic RV enrichment by DYNATE.

### 3.2 Simulation Studies: Comparing DYNATE with Competing Methods

We compared the performance of DYNATE with its competing methods: the domain-collapsing method (DC-FL and DC-SS) and the SCANG method (SCANG-B, SCANG-S, and SCANG-O). Figure 3 compares these methods’ domain-level accuracy. DYNATE and DC have similar average domain-FDP, while SCANG’s average domain-FDP is higher, sometimes above 25%. In general, DYNATE and SCANG consistently have higher a domain-sensitivity and domain-F1 score than DC. When *N*_0_ = 1, 000, DYNATE has a higher domain-sensitivity and domian-F1 score than SCANG; When *N*_0_ ≥ 2, 000, SCANG-O, SCANG-S, DYNATE-FL and DYNATE-SS have similar domain-sensitivity and domain-F1 score; all higher than SCANG-B, DC-FL, and DC-SS. The results indicate that DYNATE and SCANG have better domain-level accuracy than DC. When the sample size is small (*N*_0_ = 1, 000), DYNATE has the best domain-level accuracy.

**Figure 3:**
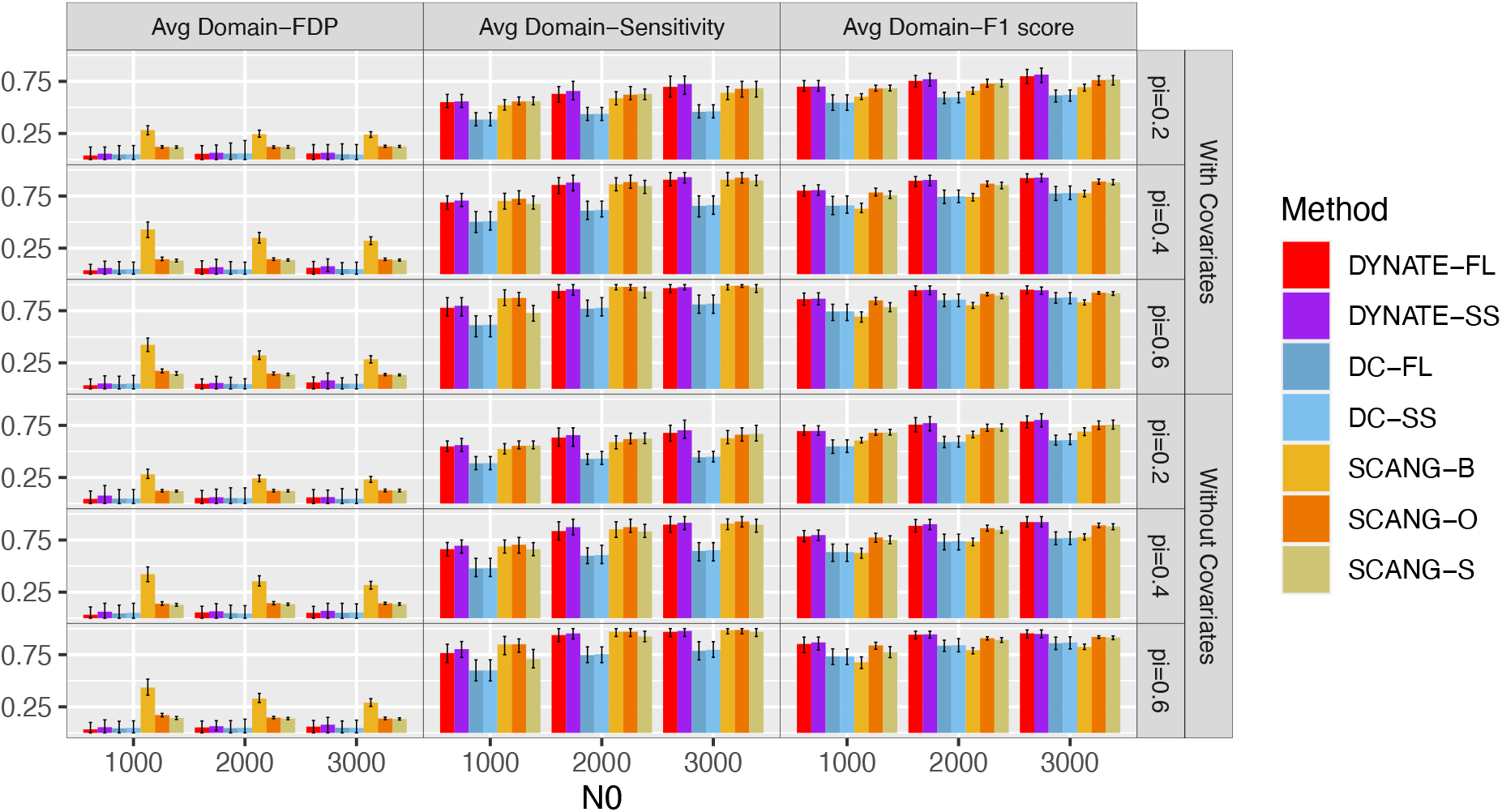
Domain-level performance comparison of the 3-layer DYNATE (DYNATE-FL, DYNATE-SS), DC (DS-FL, DC-SS) and SCANG (SCANG-B, SCANG-S, SCANG-O) under different simulation settings. The number of cases *N*_1_ = 1000; the number of controls *N*_0 ∈_ {1000, 2000, 3000}. Different colors mark different approaches of DYNATE, DC, and SCANG. The main bars mark the average performance across 100 repetitions; the error bars mark the 90% confidence intervals – the 5% and 95% quantiles over the 100 simulations. The left panel shows the average domain-FDP. The middle panel shows the average domainsensitivity. The right panel shows the average domain-F1 score.

Figure 4 compares the methods’ accuracy in pinpointing pathogenic RVs. Because DC cannot generate RV-level results, they were excluded from the comparison. SCANG, in general, has a higher RV-sensitivity than DYNATE. However, it also has extremely high average RV-FDPs, sometimes close to 1. Although the SCANG-identified regions capture more pathogenic RVs, these regions are so expansive that most of the RVs within them are neutral. This point is verified in Figure 6 for the ALS study. In contrast, DYNATE has a much lower average RV-FDP (lower than 1 *-π*). The results suggest that DYNATE is much better in identifying pathogenic-RV-enriched regions than SCANG. After we summarized RV-FDP and RV-sensitivity into an overall accuracy score, RV-F1-score, we see that DYNATE has much higher overall RV-level accuracy than SCANG. When π = 0.2, DYNATE’s average RV-level accuracy is at least four times higher than SGANG.

**Figure 4:**
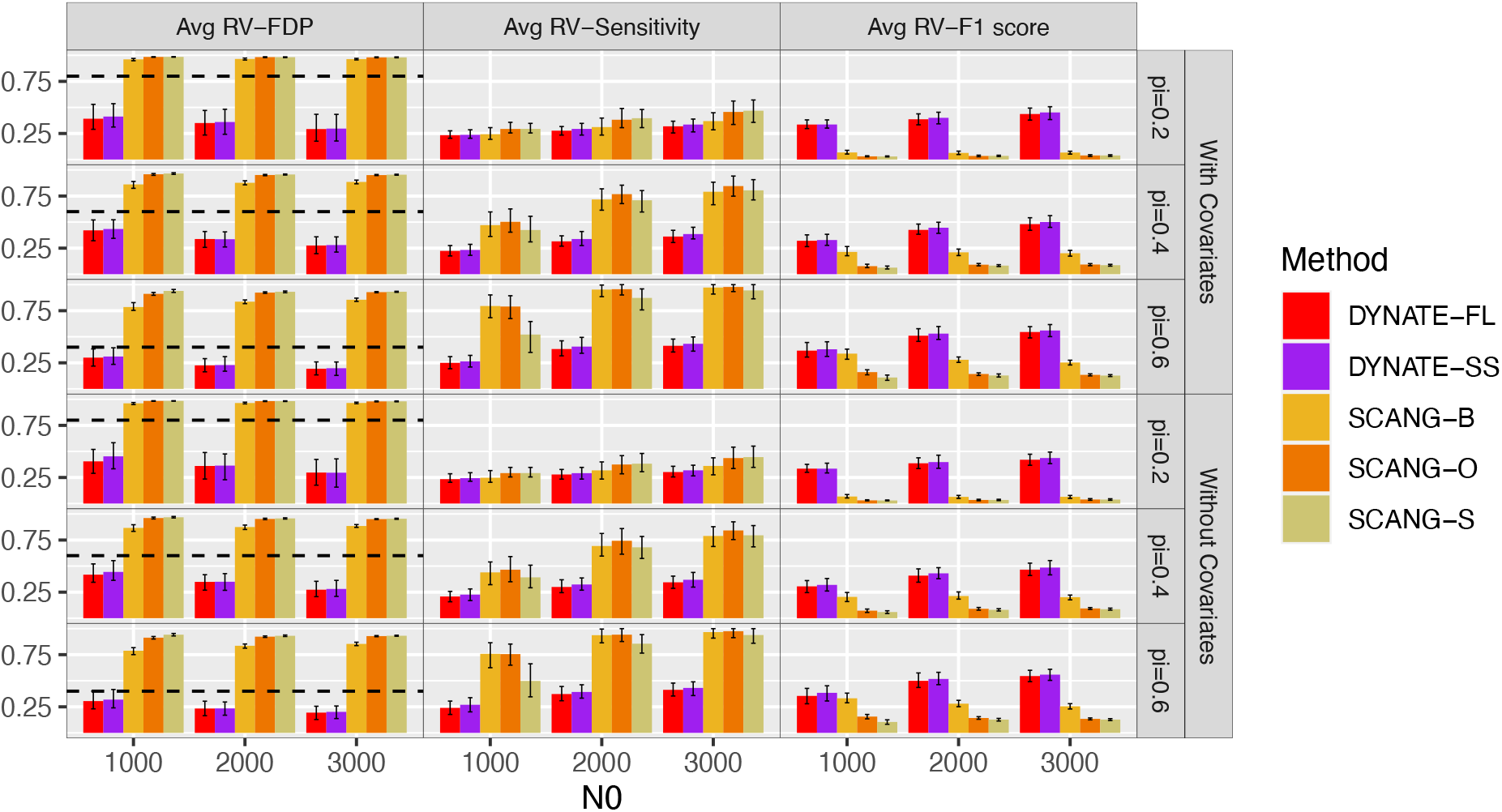
RV-level performance comparison of the 3-layer DYNATE (DYNATE-FL, DYNATE-SS) and SCANG (SCANG-B, SCANG-S, SCANG-O) under different simulation settings. The number of cases *N*_1_ = 1000; the number of controls *N*_0_ ∈ {1000, 2000, 3000}. Different colors mark different approaches of DYNATE and SCANG. The main bars mark the average performance across 100 repetitions; the error bars mark the 90% confidence intervals – the 5% and 95% quantiles over the 100 simulations. The left panel shows the average RV-FDP. The middle panel shows the average RV-sensitivity. The right panel shows the average RV-F1 score.

### 3.3 Application to an ALS Study

We applied DYNATE, DC, and SCANG to the ALS European sub-cohort, and compared their identified regions in Figure 5. All three methods identify regions overlapping with the domain *SOD1*:238186:238186_0 in *SOD1* (logOR = 2.85). *SOD1* is a well-known ALS gene(Rosen et al., 1993; Renton et al., 2014; Pansarasa et al., 2018). While SCANG also identify the regions overlapping with the *SOD1* domain, its regions are too expansive, stretching to the neighboring domains: some of them are in different genes. Also, some SCANG-identified regions extend to both pathogenic (logOR > 0) and protective (logOR *<* 0) domains, causing confusion in interpretation.

**Figure 5:**
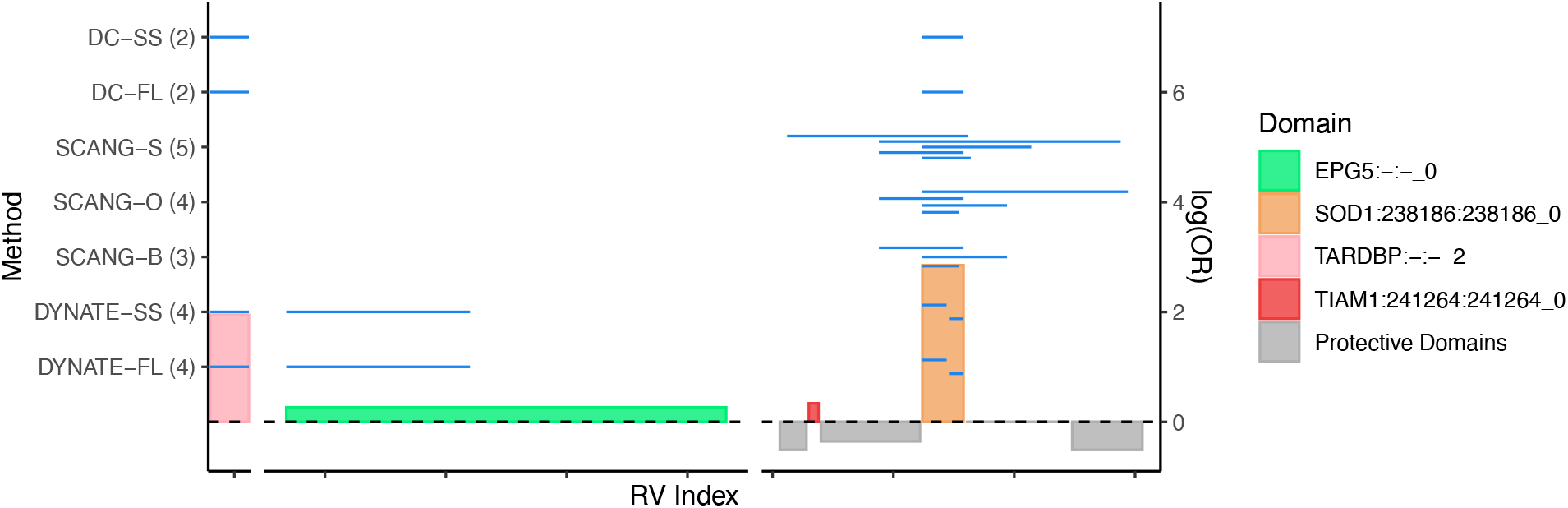
Genetic landscape of the regions identified by DYNATE, SCANG and DC. The blue solid lines mark the identified region. The regions are located in chromosome 1 (chr1), chromosome 18 (chr18) and chromosome 21 (chr21). The first row of x-axis labels the RV index in the identified regions. The RV index is ordered based on their location on each chromosome. The left y-axis labels the method that identified the region. The values in parentheses next to the labels provide a summary of the number of regions identified by each method. The right y-axis labels the domain-level logOR. Each rectangle bar represents a domain: the bar width represents the number of RVs in the domain; the bar height represents the domain-level logOR. Here, protective domains are those whose logOR < 0.

To further compare the identified regions, we compared the lengths of the DYNATE- and SCANG-identified regions (Figure 6B). Here the region length is defined as the number of RVs in the region. In general, the SCANG-identified regions are much longer than the DYNATE-identified regions: the median region lengths of DYNATE-FL and DYNATE-SS are 14; in contrast, the median region lengths of three SCANG approaches range from 36 to 46 (Supplementary Figure S2). We also compared their region-level logORs (Figure 6A). The DYNATE-identified regions have higher logORs: the median region logOR of both DYNATE-FL and DYNATE-SS are 2.22; in contrast, the median region-level logOR of three SCANG approaches range from 1.06 to 1.46. These results suggest that the DYNATE-identified regions are potentially more enriched with the pathogenic RVs than the SCANG-identified regions.

**Figure 6:**
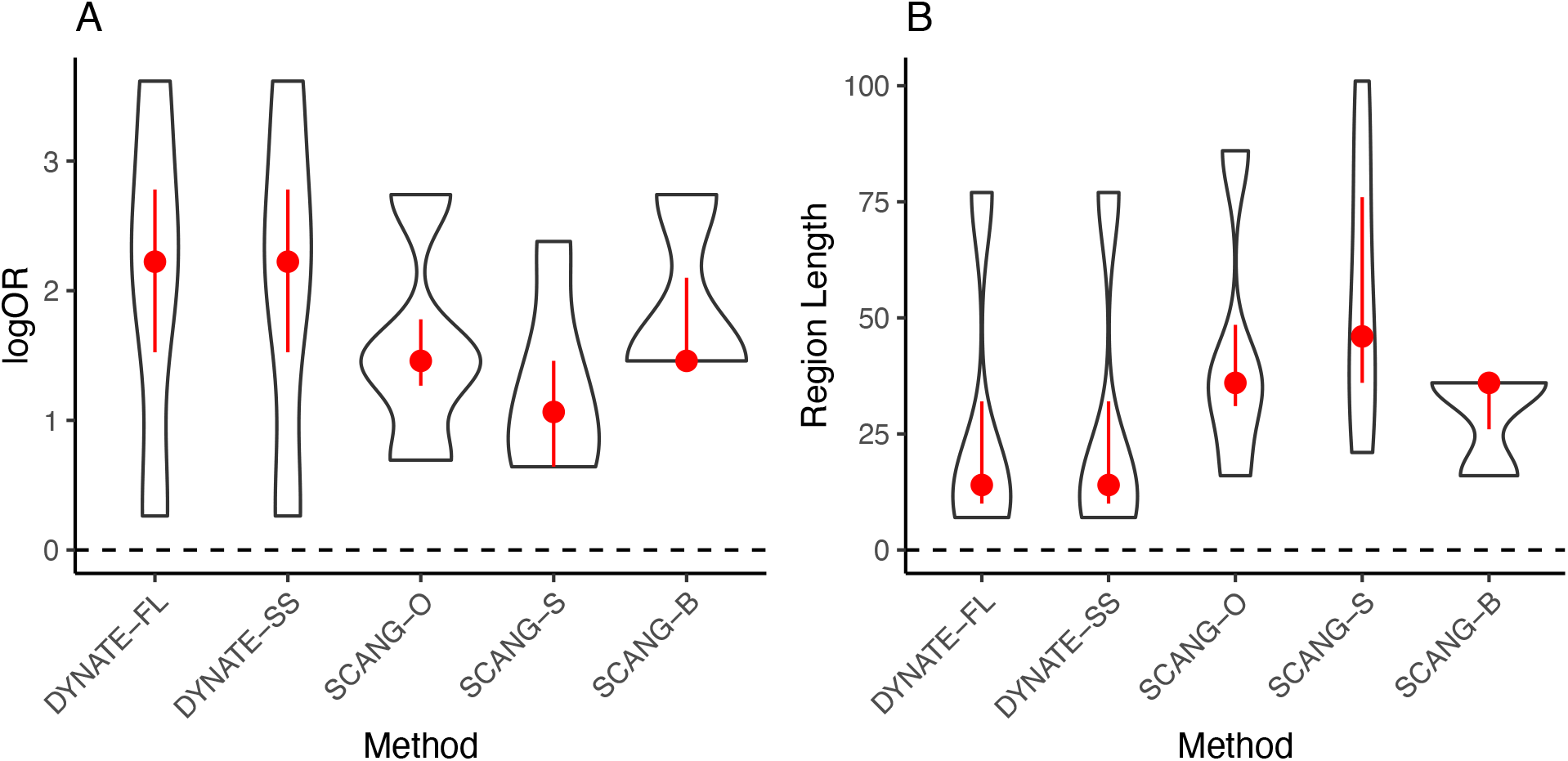
comparison of the DYNATE-identified regions and the SCANG-identfied regions. Violin plot of the distribution of detected regions’ length and logOR.The red dot indicates the median and the red vertical lines go from the 25th to the 75th percentile.

Besides the regions in *SOD1*, DC and DYNATE identified the domain *TARDBP*:–:–_2. *TARDBP* has been reported to associate with ALS in other studies (Gelfman et al., 2019; Pesiridis et al., 2009). In contrast, SCANG did not identify this domain. This domain has a slightly lower OR than *SOD1*:238186:238186_0; yet it is still high (logOR = 1.95).

In addition to the regions in *SOD1* and *TARDBP*, DYNATE identified a new region located in the *EPG5*:–:–_0. In contrast, SCANG and DC failed in identifying any regions overlapping with this domain. *EPG5* is a well-known pathogenic gene of Vici Syndrome (Cullup et al., 2013; Nam et al., 2021), another neurodegenerative disease. One previous study found that mice with the *EPG5* deficiency would exhibit key characteristics of neurodegeneration (Zhao et al., 2013). However, to the best of the authors’ knowledge, no literature has reported the association between *EPG5* and ALS on human. Thus, the domain *EPG5*:–:–_0 is an interesting finding from DYNATE and merits further investigation.

### 3.4 Computation Time

We compared the computation time of DYNATE with DC and SCANG. DC only conducts domain level analysis; thus, unsurprisingly, it is computationally efficient. Therefore, we focused on the comparison between DYNATE and SCANG. This comparison is reasonable as both dynamically scan short and long regions. We used the ALS dataset as the benchmark dataset and evaluated the computation time of both methods on a 2.10 GHz Intel Xeon Gold 6252 processor with 16 Gb of memory. DYNATE-FL took 1.7 minutes, and DYNATE-SS took 14.0 minutes. DYNATE-SS took longer because computing the score statistics with the saddlepoint-approximation is computationally demanding. Meanwhile, the SCANG package took 32.5 minutes to generate the results of SCANG-B, SCANG-S, and SCANG-O simultaneously. Most of their computation time was used to dynamically scan regions with different lengths. On average, each SCANG approach took 10.8 minutes, while each DYNATE approach took less time on average, 7.8 minutes.

Although both SCANG and DYNATE are varying-window methods, DYNATE is faster because SCANG uses an exhaustive search strategy to search all regions with lengths between the minimal and maximal constraints; in contrast, DYNATE uses a hierarchical aggregation strategy that significantly reduces the number of candidate regions (*i*.*e*., the test burden).

## 4 Discussion

This paper presents DYNATE, an innovative method formulated to detect disease-associated regions that contain RVs. Employing a dynamic aggregation and testing strategy, DYNATE examines disease associations of regions of varying lengths. As a flexible testing framework, DYNATE can be easily adapted to various studies with a range of disease outcomes, demographic and clinical covariates, statistical models, and test statistics. Below are a few additional remarks on DYNATE.

First, DYNATE is capable of extending beyond case-control studies with binary outcomes. Relying on leaf P-values, DYNATE can employ an appropriate statistical model to generate P-values when the outcome is continuous, ordinal, categorical, or censored. Thus, DYNATE is applicable for studies with various forms of outcomes.

Second, DYNATE uses a collapsing method to summarize leaf P-values. This method presumes that RVs in the leaves are associated with the phenotype in a uniform direction. However, DYNATE can also utilize other statistics like SKAT and SKAT-O, which do not impose the same directional effect on the RVs in a leaf.

Third, we advise adhering to the tuning parameter selection criteria described in Supplementary Note 2 for determining the layer number *L* and leaf size *M*. These parameters have been optimized based on our prior studies on hierarchical multiple testing (Pura et al., 2023; Li et al., 2023). While increasing layer numbers could potentially lead to more discoveries, it may also elevate the false discovery rate.

Lastly, it is important to note that DYNATE is adaptable for use with WGS data. Although this paper primarily concentrates on coding regions in WES data, largely due to the ALS study primarily using WES sequencing for most samples, it provides an effective demonstration of DYNATE’s superior performance in comparison to existing methodologies. For WGS data, broader partitions based on regulatory element maps and their potential ties to target genes in disease-relevant cell types can be considered. Moreover, nodes can be aggregated based on their similarities in functional annotation or regulatory relationships. This method presents a promising strategy for biologically informed and interpretable WGS analysis, which we aim to further explore in future studies.

## Data and Code Availability

The code to reproduce the experiments is available at https://github.com/jichunxie/DYNATE_manu. We also provide an R package to implement the proposed method, available at https://github.com/jichunxie/DYNATE. The leaf P-values in the European-based ALS corhort are available at Mendeley (DOI: 10.17632/gvykg8pwyw.1). The aggregated genotypes of case and control cohorts of the mixed population are available for download as version 3.0 of the ALS database http://www.alsdb.org.

