## Supplementary materials for "DYNATE: Localizing Rare-Variant Association Regions via Multiple Testing Embedded in an Aggregation Tree"

#### Supplementary Note 1: Approaches to derive leaf P-values

Suppose there are $N_1$ cases and $N_0$ controls in the data. The total sample size is denoted by $N = N_1 + N_0$. For subject $i$, let the outcome be $Y_k = \mathbb{I}\{\text{subject } k \text{ is a case}\}$, the non-genetic covariate be $\boldsymbol{Z}_k = (Z_{k,1}, \ldots, Z_{k,p})^T$, and the genetic vector $\boldsymbol{G}_{k,i} = (G_{k,j} : j \in \mathcal{V}(\{i\}))^T$ with $G_{k,j}$ the indicator of whether subject $k$ carries the minor allele of RV $j$. There are $D$ functional domains. Let $\mathrm{FD}_d$ be the set of qualified RVs on domain $d$, $d \in [D]$.

##### Lancaster's mid-P correction for the Fisher's exact test (FL)

First, we use the Fisher's exact test to test the association between the leaf and the disease. Then we use the Lancaster's mid-P correction to calculate the P-value. The Lancaster's mid-P correction can improve the power of the Fisher's Exact test while still controlling the type I error (Lancaster, 1961; Biddle and Morris, 2011).

Let $C_i$ denote the number of cases carrying RVs in leaf $i$, and $c_i$ be its realized value. When $H_{\{i\}}$ is null,

$$C_i \sim \mathrm{HyperGeom}(M_i, N_1, N_0),$$

where $M_i$ is the number of samples carrying RVs in leaf $\{i\}$. The p-value is given by

$$T_i = \sum_{c:P(C_i=c)<P(C_i=c_i)} \mathrm{P}(C_i = c) + \frac{1}{2} \sum_{c:P(C_i=c)=P(C_i=c_i)} \mathrm{P}(C_k = c)$$

#### Efficient score statistics with saddle point approximation (SS)

This approach calculates the collapsing score statistic based on a logistic regression model and then uses the saddle point approximation to adjust for the statistic's tail distribution (Daniels, 1954; Dey et al., 2017). The method is computationally efficient and robust when the number of subjects carrying the minor alleles is small.

Consider the logistic regression model

$$\text{logit}\{\mathrm{P}(Y_k = 1|\boldsymbol{Z}_k, \boldsymbol{G}_{k,i})\} = \gamma_0 + \boldsymbol{Z}_k^T\boldsymbol{\eta}_i + \boldsymbol{G}_{k,i}^T\boldsymbol{\beta}_i \quad \text{(S1)}$$

Here, $\boldsymbol{\eta}_i$ is the coefficient for non-genetic covariates, and $\boldsymbol{\beta}_i$ is the coefficient for the RVs on leaf $\{i\}$. To test the association between the leaf and the disease, we set the leaf hypothesis $H_{\{i\}}$ as

$$H_{\{i\},0} : \boldsymbol{\beta}_i = 0 \quad \text{versus} \quad H_{\{i\},1} : \exists\, j \in \mathcal{V}(\{i\}),\ \beta_{i,j} \neq 0.$$

We use the collapsing score test to test the hypothesis. Let $\boldsymbol{X}_i = (X_{1,i}, \ldots, X_{N,i})^T$ with $X_{k,i} = \mathbb{I}\{\sum_{j\in\mathcal{V}(\{i\})} G_{k,j} > 0\}$ be the minor allele indicator of leaf $\{i\}$. The score statistic is

$$U_i = \boldsymbol{X}_i^T(\boldsymbol{Y} - \hat{\boldsymbol{\mu}}),$$

where $\boldsymbol{Y} = (Y_1, \ldots, Y_N)^T$ is the outcome vector, and $\hat{\boldsymbol{\mu}} = (\hat{\mu}_1, \ldots, \hat{\mu}_N)^T$ is a vector with $\hat{\mu}_k$ is the estimated probablity of $Y_k$ being a case based on the null model (S1) with null $\boldsymbol{\beta}_i = 0$. The asymptotic distribution of $U_i$ when leaf $\{i\}$ is null is

$$\frac{U_i}{\sqrt{\boldsymbol{X}_i^T\hat{\boldsymbol{\Sigma}}\boldsymbol{X}_i}} \overset{asym}{\sim} N(0,1)$$

where $\hat{\boldsymbol{\Sigma}}$ is a diagonal matrix with $\hat{\boldsymbol{\Sigma}}_{k,k} = \hat{\mu}_k(1-\hat{\mu}_k)$. The calculation of P-value $T_i$ is adjusted by saddlepoint approximation to better approximate the tails of the normal distribution under the null (Dey et al., 2017).

### Supplementary Note 2: Technical details of DYNATE

#### Leaf P-value cutoff

On layer 1 (the leaf layer), let the P-value cutoff be

$$\hat{t}^{(1)} = \sup\left\{\alpha^{(1)} \le t \le \alpha : \frac{m^{(1)}t}{\{\sum_{i\in[m^{(1)}]} I(T_i \le t)\} \vee 1} \le \alpha\right\}, \tag{S2}$$

where $\alpha^{(1)} = 1/\{m^{(1)}(\log m^{(1)})^{1/2}\}$. Here, $a \vee b$ means taking the maximum of $a$ and $b$. It is easy to see that as long as $\alpha \ge 1/(\log m^{(1)})^{1/2}$, $\hat{t}^{(1)}$ must exist because $\alpha^{(1)}m^{(1)} = 1/(\log m^{(1)})^{1/2} \le \alpha$. In practice, if $\hat{t}^{(1)}$ does not exist, we set $\hat{t}^{(1)} = \alpha^{(1)}$. We reject $H_i$ if the leaf P-value $T_i < \hat{t}^{(1)}$. This testing procedure is asymptotically equivalent to the Benjamini and Hochberg procedure (Benjamini and Hochberg, 1995). Existing studies have shown that it asymptotically controls FDR under $\alpha$ (Liu, 2013; Xie and Li, 2018) when the P-values are independent or sparsely dependent (*i.e.*, a leaf P-value will not be dependent with too many other leaf P-values).

Notably, on layer 1, all leaves are either $1$-alternative or $0$-alternative. Thus, node-FDR is equivalent to the traditional FDR. In other words, the testing procedure also asymptotically controls the node-FDR.

#### Aggregation

Aggregation happens on layers $\ell$ with $\ell \ge 2$. The purpose of aggregation is to construct larger candidate regions and test for their associations.

On layer $\ell$ with $\ell \ge 2$, let $\tilde{\mathcal{B}}^{(\ell-1)} = \mathcal{B}^{(\ell-1)} \setminus \mathcal{R}$. This is a set containing all the accepted nodes on layer $\ell - 1$. Then aggregate the neighboring nodes in $\tilde{\mathcal{B}}^{(\ell-1)}$ into a new node (Figure S1). This new node is a parent node of the children nodes (being aggregated) in $\tilde{\mathcal{B}}^{(\ell-1)}$. Since domains are distinct functional and/or structural units in a protein, we do not aggregate any child nodes across different functional domains. If the last child node in a functional domain is left alone, we combine it with the preceding parent node, so that, the last parent node in the domain may contain three child nodes. After constructing all the nodes on layer $\ell$ as described, we derive the node P-values as described in (2).

For layer $\ell$, we only aggregate the accepted nodes for layer $\ell - 1$; thus, the nodes are dynamic. Accordingly, the node hypotheses are also dynamic. Testing dynamic node hypotheses is challenging because it is related to post-selection, *i.e.*, the nodes only have chances to be aggregated and tested again if they are accepted in the previous layers. Thus, we need to adaptively set the node P-value cutoff to make the tests valid and powerful.

#### Node P-value cutoffs

On higher layers, the node P-value cutoffs depend on the cutoffs on the previous layers because the nodes and the node hypotheses are dynamic.

Some nodes on layers $\ell$ ($\ell \geq 2$) may contain both null and alternative leaves. We call them mixed nodes. Denote the set of the rejected nodes by $\mathcal{R}^{(\ell)}$. Further, we denote the set of the rejected mixed nodes by $\mathcal{R}^{(\ell)}_{\text{mix}}$, the set of the rejected $100\%$-null nodes by $\mathcal{R}^{(\ell)}_{\text{null}}$, and the set of the rejected $100\%$-alternative nodes by $\mathcal{R}^{(\ell)}_{\text{altr}}$. Clearly, $\mathcal{R}^{(\ell)} = \mathcal{R}^{(\ell)}_{\text{mix}} \cup \mathcal{R}^{(\ell)}_{\text{null}} \cup \mathcal{R}^{(\ell)}_{\text{altr}}$ and the node false discoveries come from the nodes in $\mathcal{R}^{(\ell)}_{\text{mix}}$ and $\mathcal{R}^{(\ell)}_{\text{null}}$.

We derive $(\hat{t}^{(1)}, \ldots, \hat{t}^{(L)})$ recursively. The node false discovery proportion up to layer $\ell$ is

$$\text{node-FDP}^{(1:\ell)} = \frac{\sum_{\ell'=1}^{\ell} \sum_{S \in \mathcal{R}^{(\ell')}} (1 - \theta_S)}{\sum_{\ell'=1}^{\ell} |\mathcal{R}^{(\ell')}| \vee 1} = \frac{\sum_{\ell'=1}^{\ell} |\mathcal{R}^{(\ell')}_{\text{null}}|}{\sum_{\ell'=1}^{\ell} |\mathcal{R}^{(\ell')}| \vee 1} + \frac{\sum_{\ell'=1}^{\ell} \kappa^{(\ell')}}{\sum_{\ell'=1}^{\ell} |\mathcal{R}^{(\ell')}| \vee 1},$$

where

$$\text{where } \kappa^{(\ell')} = \sum_{S \in \mathcal{R}^{(\ell')}_{\text{mix}}} (1 - \theta_S).$$

Here, the superscript $(1 : \ell)$ means the quantity summarizes the results from layer 1 to layer $\ell$. Thus, for any cutoff $\{t^{(1)}, \ldots, t^{(\ell-1)}, t^{(\ell)}\}$, if we can approximate $\{(|\mathcal{R}^{(\ell)}_{\text{null}}|, \kappa^{(\ell)}) : \ell = 1, \ldots, \ell\}$, we can estimate node-FDR$^{(1:\ell)}$ $= \mathrm{E}($node-FDP$^{(1:\ell)})$. Thus, based on the previous layers' cutoffs $(\hat{t}^{(1)}, \ldots, \hat{t}^{(\ell-1)})$, we can set the P-value cutoff on layer $\ell$ as follows.

$$\hat{t}^{(\ell)} = \sup\left\{\alpha^{(\ell)} \le t^{(\ell)} \le \alpha : \frac{\sum_{\ell'=1}^{\ell} \widehat{|\mathcal{R}_{\text{null}}^{(\ell')}|}}{\sum_{\ell'=1}^{\ell} |\mathcal{R}^{(\ell')}| \vee 1} + \frac{\sum_{\ell'=1}^{\ell} \widehat{\kappa^{(\ell')}}}{\sum_{\ell'=1}^{\ell} |\mathcal{R}^{(\ell')}| \vee 1} \le \alpha\right\},$$

where $\alpha^{(\ell)} = 1/\{m^{(\ell)}(\log m^{(\ell)})^{1/2}\}$.

Now, it suffices to derive $\widehat{|\mathcal{R}_{\text{null}}^{(\ell)}|}$ and $\widehat{\kappa^{(\ell)}}$ for any layer $\ell$.

**To derive $\widehat{|\mathcal{R}_{\mathbf{null}}^{(\ell)}|}$.** Under the ideal case, the null P-values should asymptotically follow uniform distributions. Unfortunately, the null distributions of the node P-values might change because of the post-selection process in aggregation. However, when most nodes are $100\%$-null, the null p-values should be asymptotically uniformly distributed (Li et al., 2023). Thus, the post-selection process will not affect the null distributions too much. Therefore, we expect $|\mathcal{R}_{\text{null}}^{(\ell)}|$ to be close to or smaller than $tm^{(\ell)}$, where $m^{(\ell)} = |\mathcal{B}^{(\ell)}|$ is the total number of valid nodes on layer $\ell$.

**To derive $\widehat{\kappa^{(\ell)}}$.** We use $\mathrm{E}(\kappa^{(\ell)}|\mathcal{X}^{(\ell-1)})$ to approximate $\kappa^{(\ell)}$, where $\mathcal{X}^{(\ell-1)}$ is the testing path up to layer $\ell-1$. Note that

$$\mathrm{E}(\kappa^{(\ell)} \mid \mathcal{X}^{(\ell-1)}) = \sum_{d_1,d_2} w_{d_1,d_2}^{(\ell)}\, \mathrm{E}(\kappa_{d_1,d_2} \mid \mathcal{X}^{(\ell-1)}),$$

where $w_{d_1,d_2}^{(\ell)}$ is the proportions of the mixed nodes containing $d_1$ alternative and $d_2$ null leaves among all rejected mixed nodes; $\mathrm{E}(\kappa_{d_1,d_2})$ is the expected false discoveries among them. It suffices to approximate $w_{d_1,d_2}^{(\ell)}$ and $\mathrm{E}(\kappa_{d_1,d_2} \mid \mathcal{X}^{(\ell-1)})$. On layer 1, we call the leaves with the smallest $\lfloor\sqrt{m^{(1)}}\rfloor$ P-values as pseudo-alternative leaves and the rest as pseudo-null leaves.

- To approximate $w_{d_1,d_2}^{(\ell)}$: We classify rejected nodes in $\mathcal{R}^{(\ell)}$ (given a candidate cutoff $t$) into groups: each group contains nodes with the same numbers of pseudo-null and pseudo-alternative leaves. Then, $w_{d_1,d_2}^{(\ell)}$ is approximated by the proportion of nodes with $d_1$ pseudo-alternative and $d_2$ pseudo-null leaves.
- To approximate $\mathrm{E}(\kappa_{d_1,d_2} \mid \mathcal{X}^{(\ell-1)})$: For fast computation, we randomly sample $B$ mixed

nodes, each with $d_1$ pseudo-alternative and $d_2$ pseudo-null leaves. We calculate node P-values for those nodes. Given the P-value cutoff $t$, we will get the rejected sets, denoted by $\tilde{\mathcal{R}}_{d_1,d_2}$. We then approximate $\mathrm{E}(\kappa_{d_1,d_2} \mid \mathcal{X}^{(\ell-1)})$ by

$$\hat{\kappa}_{d_1,d_2} = \frac{d_2}{d_1 + d_2} \frac{|\tilde{\mathcal{R}}_{d_1,d_2}|}{B}. \quad \text{(S3)}$$

Because the rejected leaves only take a small proportion among all pseudo-null and pseudo-alternative leaves, such approximation is accurate enough to perform reliable testing. Also, for any possible $(d_1, d_2)$, $|\tilde{\mathcal{R}}_{d_1,d_2}|$ only needs to be calculated once. We may calculate them before the hierarchical aggregation algorithm begins. This saves significant computational time.

#### Tuning parameter selection

The layer number $L$ and leaf size $M$ are tuning parameters. Here we discuss how to set these parameters.

- We first set $M_0$ as the smallest value so that we have the power to identify at least one leaf on layer 1. To ensure the stability of the statistical testing, we suggest $M_0 \geq 7$. Denote the number of RVs in domain $d$ by $V_d$. Most qualified nodes are formed by aggregating two child nodes on the previous layer; thus, a null domain $d$ will likely remain till the layer $\lceil \log_2(V_d/M_0) \rceil$. Since most of the domains are null domains, to ensure 50 nodes on layer $L$, we set $L = \lceil \log_2(V'/M_0) \rceil$, where $V'$ ranks the 50 largest among all $\{V_d : d \in [D]\}$.
- We set the leaf size $M$ as the maximum value to keep $L$ unchanged.

$$M = \max\{M' : \lceil \log_2(V'/M') \rceil = L\}.$$

#### Supplementary Figures

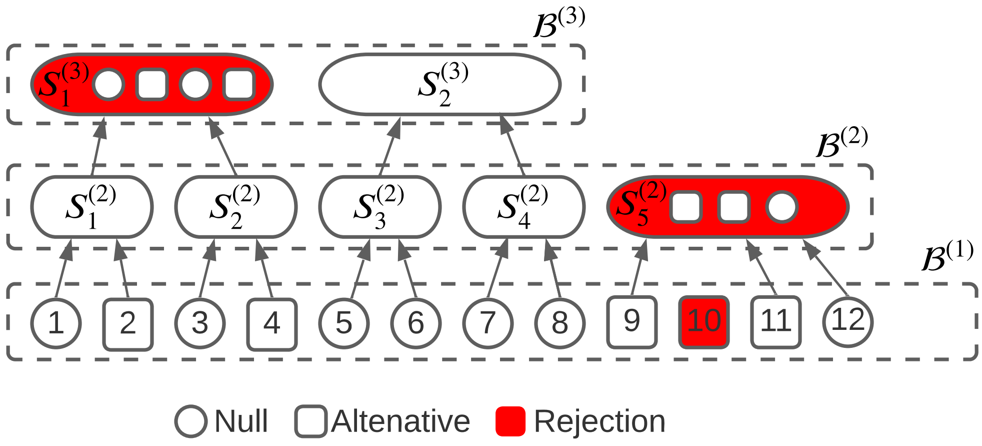


Figure S1: A toy example of a three-layer DYNATE analysis. Node $i$ on layer $\ell$ is denoted by $S_i^{(\ell)}$. On layer 1, we reject leaf $\{10\}$. On layer 2, we aggregate the neighboring accepted leaves into five nodes, from which we reject $S_5^{(2)} = \{9, 11, 12\}$. On layer 3, we reject $S_1^{(3)}$. Clearly, $S_{10}^{(1)}$ is $1$-alternative, $S_5^{(2)}$ is $2/3$-alternative, and $S_1^{(3)}$ is $1/2$-alternative. The empirical node-FDR of this example is $5/18$.

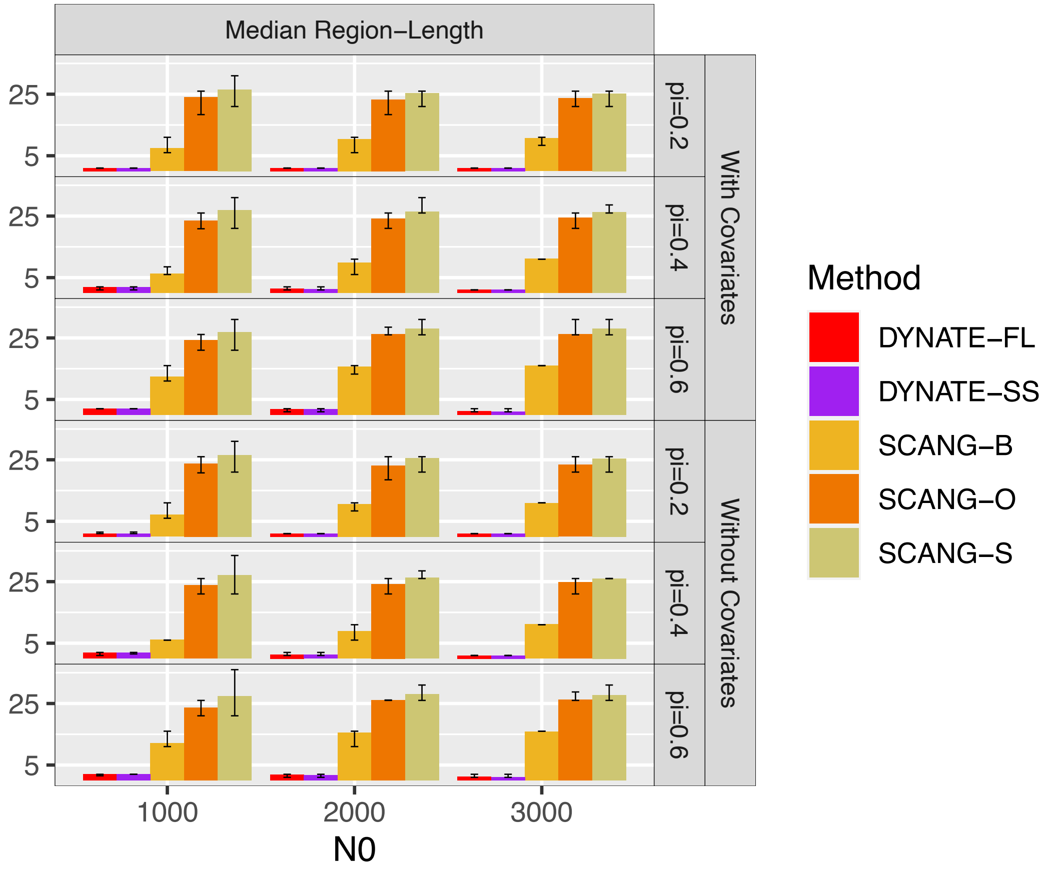


Figure S2: The median length of the DYNATE- and SCANG-identified regions in the ALS European sub-cohort data. The error bars in the bar plot are the $90\%$ confidence intervals, which are calculated as the $5\%$ and $95\%$ quantiles over the $100$ simulations.
